# Androgens enhance adult hippocampal neurogenesis in males but not females in an age-dependent manner

**DOI:** 10.1101/539296

**Authors:** Paula Duarte-Guterman, Dwayne K. Hamson, Stephanie Lieblich, Steven R. Wainwright, Carmen Chow, Jessica Chaiton, Neil V. Watson, Liisa A.M. Galea

## Abstract

Androgens (testosterone and dihydrotestosterone) increase adult hippocampal neurogenesis by increasing survival of new neurons in male rats and mice via an androgen receptor pathway, but it is not known whether androgens regulate neurogenesis in females and whether the effect is age-dependent. We investigated the effects of dihydrotestosterone (DHT), a potent androgen, on neurogenesis in young adult and middle-aged males and females. Rats were gonadectomized and injected with the DNA synthesis marker, bromodeoxyuridine (BrdU). The following day rats began receiving daily injections of oil or DHT for 30 days. We evaluated cell proliferation (Ki67) and survival of new neurons (BrdU and BrdU/NeuN) in the hippocampus of male and female rats using immunohistochemistry. As expected, DHT increased the number of BrdU+ cells in young males but surprisingly not in middle-aged male rats or in young and middle-aged females. In middle age, DHT increased the proportion of BrdU/NeuN cells, an effect driven by females. AR expression also increased with aging in both female and male rats, which may contribute to a lack of DHT neurogenic effect in middle age. Our results indicate that DHT regulates adult hippocampal neurogenesis in a sex- and age-dependent manner.

## Main text

Neurogenesis, the production of new neurons, in the hippocampus continues through the life span of most mammals studied to date (1). Sex hormones (estrogens and androgens) regulate different aspects of hippocampal neurogenesis: e.g. proliferation and/or survival of these new neurons in rodents (reviewed in (2,3)). There is also evidence of sex differences in how hormones regulate neurogenesis. For example, estradiol regulates cell proliferation and survival of new neurons in female but not male rats (4). We have previously shown that androgens (testosterone and dihydrotestosterone) increase the survival of new neurons but not cell proliferation in the hippocampus of male rats and mice via an androgen receptor (AR) pathway (5,6,7). However it is not known whether androgens regulate any aspects of adult neurogenesis in females, even though ARs are expressed in the female hippocampus (8). In addition to sex, age can also modulate the effects of hormones on hippocampal neurogenesis. In middle age, neurogenesis decreases (9) while reduction of corticosteroid levels (by adrenalectomy) (10) and exercise (11) restore neurogenesis levels in aged rodents. With aging, the hippocampus also loses its ability to respond to estrogens in female rats (12,13). For instance, estradiol increases cell proliferation in the hippocampus in young but not middle-aged nulliparous female rats (12,13). The objective of this study was to investigate the effects of dihydrotestosterone (DHT) on hippocampal neurogenesis (proliferation and survival of new neurons) in young and middle-aged male and female rats.

At 2 months (∼70 days old, young) and 11-12 months of age (middle-aged), male and female Sprague–Dawley rats were gonadectomized and allowed to recover for one week (n=5-8/group) (4–6,14). One week allows for circulating gonadal hormone levels to decrease to very low or undetectable levels (4,5,15,16). One day after ovariectomy, estradiol levels are undetectable in female rats (15) and approximately 5 days after gonadectomy circulating estradiol and testosterone decrease to approximately 10% of their original levels or to undetectable levels in males (17,18). We chose 11-12 months as middle-age as rats can live up to ∼24 months, at 12 months the levels of neurogenesis are substantially decreased compared to young adults (9,19–21), and at 12 months sexual motivation and fecundity is significantly reduced in both sexes (22–24). After the one week recovery period, all animals received a single intraperitoneal injection of bromodeoxyuridine (BrdU; 200 mg/kg) to label dividing cells and their progeny (6). The following day, chronic hormone or vehicle treatment began. Males and females were injected subcutaneously with either 0.25 mg dihydrotestosterone (DHT in 0.1 ml of sesame oil) or an equivalent volume of sesame oil for 30 days. The dose of DHT chosen in this study was the lowest dose examined that increased neurogenesis in castrated young adult male rats (5,6). Twenty-four hours after the final injection, animals were overdosed with sodium pentobarbitol, and perfused with 4% paraformaldehyde, then brains were collected, sectioned using a freezing microtome and processed for BrdU (survival of 30 day old cells), Ki67 (cell proliferation marker), androgen receptor (AR), and colabelled for BrdU/NeuN (new neurons using NeuN, marker for mature neurons) immunohistochemistry. The following primary antibodies were used with diaminobenzidine (DAB) chromogen: mouse anti-BrdU monoclonal (1:200; Roche Cat#11170376001 (25)), rabbit anti-Ki67 polyclonal (1:3000; Vector Laboratories Cat# VP-K451 (26)), and rabbit anti-androgen receptor monoclonal (1:100; Abcam Cat# ab133273 (27)). For fluorescence double labelling the following primary antibodies were used: rat anti-BrdU (1:500; Bio-Rad/ABD Serotec Cat#OBT0030S (28)) and mouse anti-NeuN (1:250; Millipore Cat# MAB377 (29)). Detailed protocols are found in (6,7). Thus, in this experiment, BrdU+ cells were 30 day-old daughter cells from progenitor cells that had been synthesizing DNA for a 2-hour period 31 days before euthanasia. A subset of samples were used to measure DHT levels in serum collected on the day of perfusion (stored at −20°C) using a commercial ELISA kit (IBL-America, Cat#IB59116 (30)). All samples were run in duplicate following the manufacturer’s protocol. The DHT antibody is highly specific with 8.7% cross-reactivity with testosterone and 0.2% cross-reactivity with androstenedione; the sensitivity is 6.0 pg/ml. Average intra-assay coefficients of variation were <15%. All protocols were approved by the Animal Care Committee at the University of British Columbia and conformed to the guidelines set out by the Canadian Council on Animal Care.

A researcher blind to experimental conditions counted all BrdU+ and Ki67+ cells in both hemispheres for each section for the entire rostrocaudal extent of the granule cell layer (GCL) including the sub granular zone, defined as the 50 µm band between the GCL and the hilus (13-15 sections in total per animal). We used a modified optical fractionator method (31,32) to estimate the total number of BrdU+ and Ki67+ cells in the GCL and hilus, as has been used before (5,7,33–37). Total cells were calculated by multiplying the total number of cells counted by 10 to account for the fact that we used 1/10 series of sections for each immunohistochemistry procedure. GCL and hilus volumes were quantified from digitized images using Cavelier’s principle, multiplying the sum of the area of each section by the section thickness (40 µm) (38). Densities of BrdU+ and Ki67+ were calculated by dividing the total number of cells by the GCL volume. Densities (total cells per unit volume) were used as there are sex differences in the volume of the dentate gyrus in rats (39,40). We assessed expression of AR by quantitative densiometric analysis using ImageJ (U. S. National Institutes of Health, Bethesda, Maryland, USA, http://imagej.nih.gov/ij/). Photomicrographs of four hippocampal sections per animal were taken at 40X magnification using the same exposure and gain settings. Optical density (OD) was assessed in the CA1, CA3, and GCL regions by placing six circles (40 µm diameter) along these regions. OD levels were corrected for background levels using areas with no immunoreactivity (i.e. molecular layer of the dentate gyrus and radiatum layer of the hippocampus). To determine whether BrdU+ cells were of a neuronal phenotype, in a subset of brains, BrdU+ cells were examined for co-labeling with NeuN (neuronal marker) for 50 cells (young adult group) or all cells (middle-aged group). All analyses were performed using Statistica v.8.0 (StatSoft Inc, Tulsa, OK). Volume of the dentate gyrus (GCL and hilus) and density of BrdU+ cells were analyzed using repeated measures analysis of variance (ANOVA) with age (young, middle-aged), sex (male, female), and treatment (DHT, oil) as between subjects factors, and region (GCL, hilus) as a within subjects factor. AR OD was analyzed using repeated measures ANOVA with age (young, middle-aged), sex (male, female), and treatment (DHT, oil) as between subjects factors, and region (CA1, CA3, GCL) as a within subjects. Density of Ki67+ cells, proportion of BrdU/NeuN co-labeled cells, and serum DHT concentrations were analyzed using ANOVA with age (young, middle-aged), sex (male, female), and treatment (DHT, oil) as between subjects factors. When appropriate, post-hoc analysis used the Neuman-Keul’s procedure. Test statistics were considered significant if p ≤ 0.05.

Regardless of age and treatment, the volume of the GCL and hilus were larger in males compared to females (main effect of sex; F(1,42)=4.42; P<0.05; Table 1). The volume of the GCL and hilus increased with age irrespective of sex and treatment (main effect of age; F(1,42)=47.37; P<0.0001). In the case of the hilus, the volume increased with aging more so in males than in females (interaction between region, age, and sex; F(1,42)=20.20; P<0.0001). As expected, the volume of the hilus was larger than the volume of the GCL (main effect of region; F(1,44)=1450.14; P<0.0001). Treatment with DHT did not significantly affect GCL or hilus volumes (all P’s>0.08). To account for the sex differences in GCL and hilus volumes, we present BrdU+ and Ki67+ cell counts as densities (total cells per unit volume).

**Table 1.** Mean (SEM) volume of the granule cell layer (GCL) and the hilus (mm^3^) and BrdU density in the hilus in gonadectomized young and middle-aged male and female rats treated with oil or dihydrotestosterone (DHT). The volume of the hilus was larger than the volume of the GCL (main effect of region; P<0.0001). Males had a larger volume of GCL and hilus than females (main effect of sex; P<0.05). The volume of the GCL and hilus increased with age (main effect of age; P<0.0001) and in the hilus, the volume increased with aging more so in males than females (interaction between region, age, and sex; P<0.0001). DHT treatment did not affect the volume of the GCL and hilus (all P’s>0.08). The density of BrdU+ cells was not significantly affected by age, sex or treatment (all P’s>0.9). Sample size is number of individuals.

| Age | Sex and Treatment | Sample size | GCL volume (mm<sup>3</sup>) | Hilus volume (mm<sup>3</sup>) | Hilus BrdU density (per mm<sup>3</sup>) |
| --- | --- | --- | --- | --- | --- |
| Young | *Females* |  |  |  |  |
|  | Oil | 7 | 2.51 (0.12) | 6.09 (0.19) | 85.14 (15.34) |
|  | DHT | 8 | 2.46 (0.08) | 6.00 (0.28) | 76.06 (7.06) |
|  | *Males* |  |  |  |  |
|  | Oil | 5 | 2.31 (0.19) | 5.32 (0.49) | 96.18 (23.29) |
|  | DHT | 6 | 2.62 (0.21) | 5.37 (0.51) | 134.70 (17.38) |
| Middle Age | *Females* |  |  |  |  |
|  | Oil | 6 | 2.33 (0.09) | 7.67 (0.42) | 20.69 (7.89) |
|  | DHT | 6 | 2.45 (0.15) | 7.33 (0.28) | 6.40 (2.49) |
|  | *Males* |  |  |  |  |
|  | Oil | 6 | 2.94 (0.25) | 9.05 (0.79) | 6.06 (1.41) |
|  | DHT | 6 | 2.87 (0.22) | 10.05 (0.60) | 4.65 (1.63) |

As expected, DHT treatment increased serum levels of DHT in males and females regardless of age (main effect of treatment; F(1,17)=36.67; P<0.0001). The mean (±SEM) DHT level in oil treated rats was 38.55 ± 2.99 pg/ml and in DHT-treated rats was 73.64 ± 8.70 pg/ml. DHT levels were significantly higher in females than males regardless of treatment (main effect of sex; F(1,17)=26.6; P<0.0001). The mean (±SEM) DHT level in DHT-treated females was 69.58 ± 8.66 pg/ml and in DHT-treated males was 40.02 ± 3.79 pg/ml. No significant differences were detected in DHT levels with age (P’s>0.3).

DHT treatment increased the density of BrdU+ cells in the GCL in young males (P<0.001) but not in young females or middle-aged rats of both sexes (all P’s>0.9; interaction between age, sex, treatment and region; F(1,42)=4.03; P=0.05; Figure 1). The density of BrdU+ cells in the GCL was significantly higher in the young compared to the middle-aged rats irrespective of sex and treatment, as expected (main effect of age; F(1,42)=647.85; P<0.0001). In the hilus, the density of BrdU+ cells was not significantly affected by age, sex or treatment (all P’s>0.9; Table 1). To determine how many BrdU+ cells were neurons, we examined the colabelling of BrdU and NeuN (a mature neuronal marker; BrdU/NeuN) in the GCL. Aging decreased the proportion of BrdU/NeuN colabeled cells in the dentate gyrus (main effects of age; F(1,39)=27.98; P<0.0001). In the young group, sex and treatment did not affect the percent of BrdU/NeuN colabeled cells (all P’s>0.18; Table 2). In middle age, DHT treatment increased the proportion of BrdU/NeuN cells (interaction between age and treatment; F(1,39)=4.77; P=0.03), and this effect was driven by the middle-aged females (P=0.005) compared to the middle-aged males (P=0.48).

**Figure 1.**
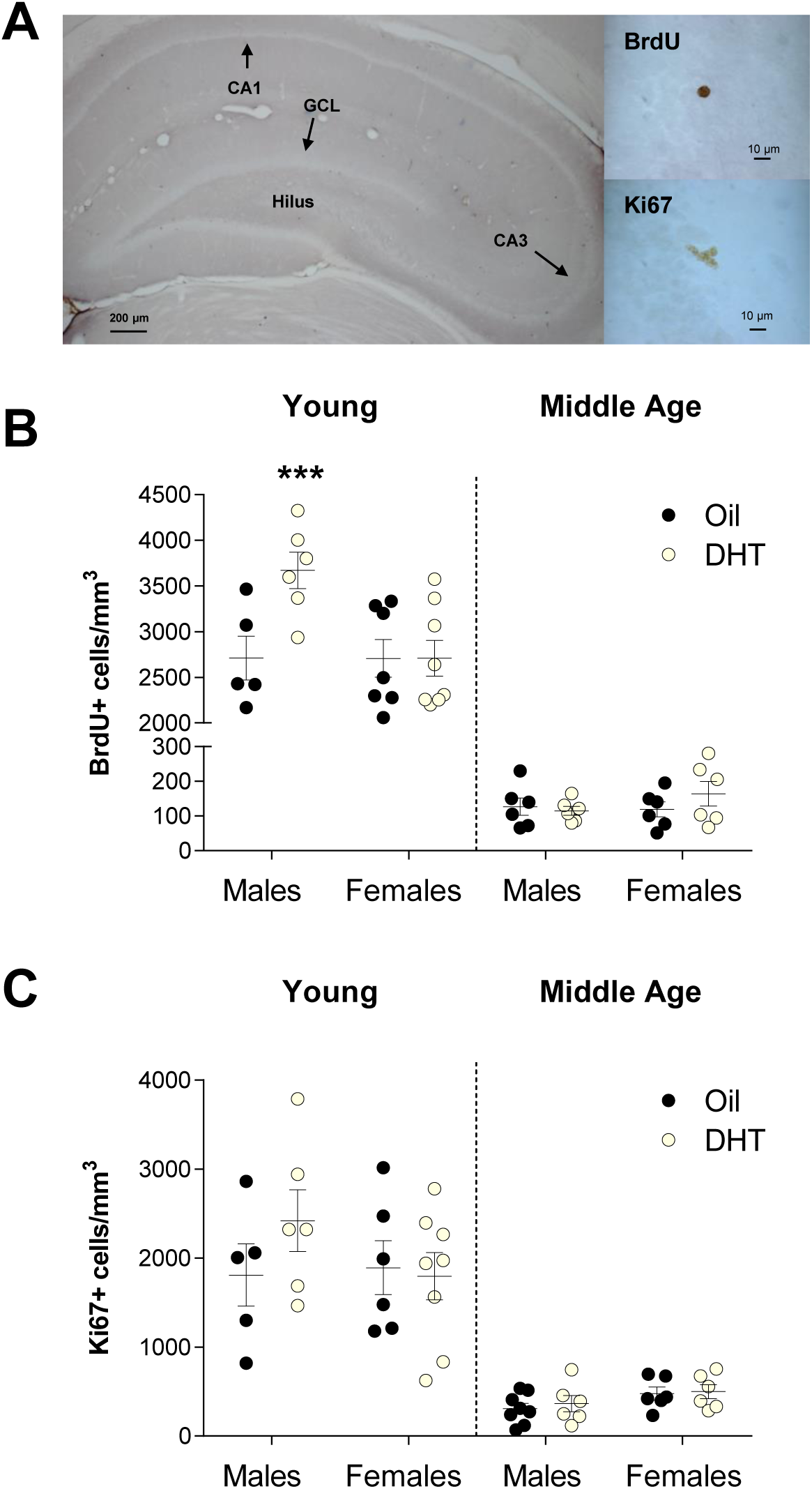
Chronic dihydrotestosterone (DHT) treatment increases the density of BrdU+ cells in the dentate gyrus of young male adult rats but has no effects in middle-aged males and young and middle-aged female rats. (**A**) Photomicrographs of a representative section of dentate gyrus with the granule cell layer (GCL), hilus, CA1 and CA3 regions and representative BrdU+ and Ki67+ cells in the subgranular zone. (**B**) Mean ± SEM total number of BrdU+ cells in the granule cell layer of the dentate gyrus in young and middle-aged male and female rats. In young males, DHT increased the density of BrdU+ cells relative to the oil treatment group (*** P < 0.001; interaction between age, sex, treatment, and region; P=0.05). (**C**) Mean ± SEM total number of Ki67+ cells in the granule cell layer of the dentate gyrus in young and middle-aged male and female rats. Chronic DHT treatment did not affect cell proliferation in the hippocampus of young or middle-aged male and female rats. Circles represent individual data points (number of individuals).

**Table 2.** Mean (SEM) percentage of cells co-expressing BrdU and NeuN in the GCL in gonadectomized young and middle-aged male and female rats treated with oil or dihydrotestosterone (DHT). The proportion of BrdU+ cells colabelled with NeuN was significantly higher in young compared to middle-aged animals (main effects of age; P<0.0001. In middle age, but not in young adults, DHT increased the proportion of BrdU/NeuN colabelled cells relative to the oil treatment group (interaction between age and treatment) but this effect was driven by the middle-aged females (P=0.005) compared to the middle-aged males (P=0.48). ** P<0.01 relative to oil treatment group. Sample size is number of individuals.

| Age | Sex and Treatment | Sample size | BrdU/NeuN % |
| --- | --- | --- | --- |
| Young | *Females* |  |  |
|  | Oil | 5 | 87.89 (2.27) |
|  | DHT | 8 | 87.16 (1.10) |
|  | *Males* |  |  |
|  | Oil | 4 | 88.68 (1.94) |
|  | DHT | 5 | 84.24 (3.12) |
| Middle Age | *Females* |  |  |
|  | Oil | 6 | 63.80 (3.43) |
|  | DHT | 6 | 81.32 (5.82)** |
|  | *Males* |  |  |
|  | Oil | 6 | 67.03 (5.39) |
|  | DHT | 6 | 70.88 (7.00) |

DHT treatment did not affect the density of Ki67+ cells in young or middle-aged males and females (all P’s>0.25; Figure 1). However, as expected, there were more Ki67+ cells in the GCL of the young compared to middle-aged rats (main effect of age; F(1,43)=97.18; P<0.0001) irrespective of sex and treatment.

Finally, we performed a qualitative and quantitative (OD) analysis of AR expression in the hippocampus and both methods obtained similar results (Table 3; Figure 2, respectively). AR OD increased significantly with aging in the CA1, CA3, and GCL, regardless of sex, and DHT treatment increased the expression of AR in the CA1 in middle-aged rats of both sexes (interaction between region, age and treatment; F(2,48)=8.1; P<0.001; Figure 2). However, DHT treatment increased AR OD in the CA1, regardless of age, more so in females (P=0.0001) than in males (P=0.056; interaction between region, sex and treatment; F(2,48)=3.3; P<0.05; Figure 2). Qualitatively, in oil treated animals, AR-ir cells were absent throughout the GCL in young males and female rats but were expressed at low levels in middle-aged male and female rats. In the CA3, we found low to absent levels of AR-ir cells in young rats of both sexes and expression increased in middle-aged rats. In the CA1, we found low to intermediate levels of AR-ir in oil treated young rats and levels increased with aging and DHT treatment in both sexes (Figure 2).

**Figure 2.**
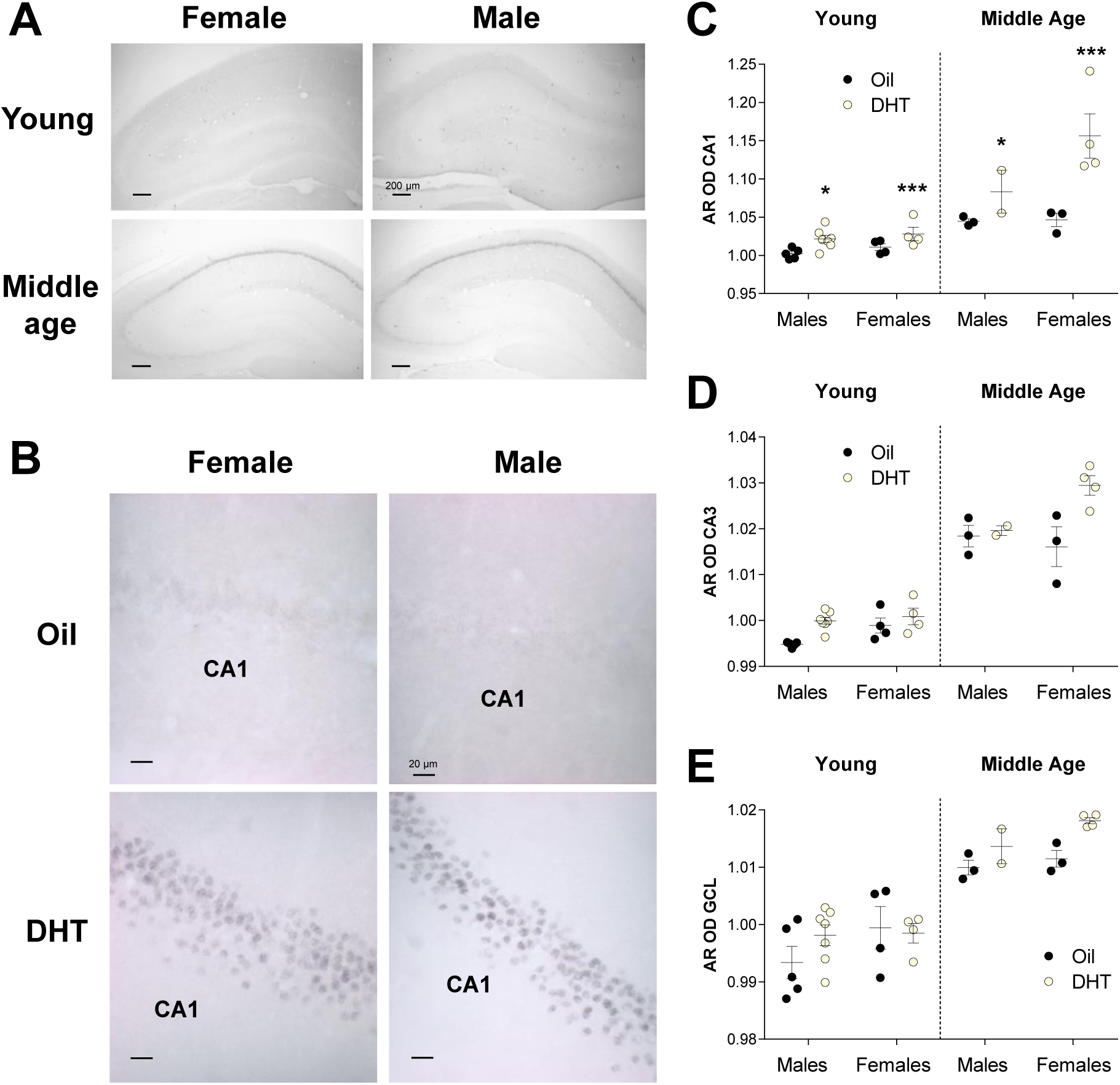
Chronic dihydrotestosterone (DHT) treatment increases AR optical density in the CA1 region of the hippocampus in young and middle-aged male and female rats. (**A**) Representative images of androgen receptor (AR) expression in the hippocampus of young and middle-aged gonadectomized oil treated female and male rats. AR expression increased with aging in the CA1, CA3 and GCL in both sexes. (**B**) Representative images of androgen receptor (AR) expression in the CA1 region of the hippocampus in young male and female rats treated with oil or DHT. AR is lowly expressed in the CA1 of gonadectomized oil treated animals. DHT increases the expression of AR in both female and male rats. Optical density (OD) for AR (mean ±SEM) was measured in the CA1 (**C**), CA3 (**D**), and GCL (**E**) regions. AR OD increased significantly with aging in the CA1, CA3, and GCL, regardless of sex, and DHT treatment increased the expression of AR in the CA1 in young and middle-aged rats of both sexes (interaction between region, age and treatment; P<0.001). DHT treatment increased AR OD in the CA1, regardless of age, more so in females than in males (interaction between region, sex and treatment; P<0.05). Asterisks denote significant differences between oil and DHT treatment (*P≤0.05; ***P<0.001). Circles represent individual data points (number of individuals).

**Table 3.** Androgen receptor (AR) expression in the hippocampus using a relative rating scaling: absent (0), light (+), intermediate (++), and robust (+++) in the granule cell layer (GCL), CA1, and CA3 regions in young and middle-aged male and female rats treated with oil or dihydrotestosterone (DHT).

| Age | Sex and Treatment | AR expression |  |  |
| --- | --- | --- | --- | --- |
|  |  | GCL | CA1 | CA3 |
| **Young** | *Females* |  |  |  |
|  | Oil | 0 | + | 0/+ |
|  | DHT | 0 | ++/+++ | 0 |
|  | *Males* |  |  |  |
|  | Oil | 0 | 0/+ | 0 |
|  | DHT | 0 | ++ | 0 |
| **Middle Age** | *Females* |  |  |  |
|  | Oil | 0/+ | ++ | + |
|  | DHT | + | +++ | +/++ |
|  | *Males* |  |  |  |
|  | Oil | 0/+ | ++ | + |
|  | DHT | + | ++/+++ | + |

In the present study, we found that chronic (30 days) DHT increased survival of new neurons but not cell proliferation in the hippocampus of gonadectomized young adult male rats, consistent with our previous research in male rats and mice (5–7). This is also in line with previous work showing that castration decreases survival of new neurons 24-30 days after BrdU injection but has no effect on cell proliferation (5,41). Shorter testosterone treatment (3, 15 or 21 days) has no effect on hippocampal neurogenesis (42–45) indicating that a longer exposure (30 days) to androgens is required to increase neurogenesis in the hippocampus at least in physiological doses as higher doses can decrease neurogenesis (46,47). Thus, collectively while longer term exposure to androgens increases survival of new neurons in the dentate gyrus, androgens do not appear to influence cell proliferation in male rats (this study and (5,6)), mice (7), or voles (48).

In contrast to young adult males, DHT did not affect survival of new neurons or cell proliferation in gonadectomized young adult female rats. We previously found that estradiol modulates cell proliferation and survival of new neurons in young adult female rats (4,49–52) but has no significant effect on neurogenesis in adult male rats (4,5). We found that females had higher serum DHT levels than males regardless of treatment or age. The higher DHT levels in females were likely due to the dose being somewhat higher in females compared to males. So it is possible that these higher circulating DHT levels resulted in an eliminated response to neurogenesis, although we observed an increase in cell fate (BrdU/NeuN) with DHT in middle-aged females (discussed below). Together our results suggest that sex steroids have sex-specific effects on hippocampal neurogenesis with androgens modulating neurogenesis in young adult males and estrogens modulating neurogenesis in young adult females. While these findings may not seem surprising, it is important to understand that both sexes have ARs and estrogen receptors (ERs) but these receptors are responding to respective hormones in a sex-specific way to modulate neurogenesis in the hippocampus. Both ERα and ERβ are expressed in the hippocampus (CA1, CA3, GCL) in males and females and no sex differences exist in their expression (53,54). Intriguingly, there are more ERs in the GCL than there are ARs in both sexes, and ERs have been detected on proliferating cells (Ki67+ or BrdU+ cells) or immature neurons (doublecortin expressing cells) in adult male and female rats (50,55,56) but to our knowledge ARs have not been detected on proliferating cells or immature neurons in the dentate gyrus in male rats and mice (6,7). Our findings are unlikely to involve ERs (α or β) due to the use of DHT in this study. DHT is a non-aromatizable androgen that binds with high affinity to AR. However, 5α-androstane-3β, 17β-diol (3β-Adiol), a DHT metabolite, has been shown to function as an ERβ ligand (57). In order to avoid any possible ERβ activation, we used the lowest possible dose of DHT that increases neurogenesis in young males (5). From previous work, estradiol does not increase neurogenesis in males and decreases neurogenesis (survival of new neurons) in females (4). In addition, in young male rats, the same dose of DHT regulates neurogenesis via ARs as blocking AR with flutamide eliminates the DHT-induced increase in new neuron survival (6). Together this suggests that the DHT regulation of hippocampal neurogenesis is mediated by the AR.

Perhaps surprisingly, the effect of DHT to modulate survival of new neurons was absent in middle-aged males and females. This effect is consistent with the recent work of Moser et al. (58) who found that testosterone did not affect the number of immature neurons (using doublecortin), in middle-aged (13 months) or aged (23 months) male rats. This would suggest that with aging, the dentate gyrus loses its ability to respond to sex steroids. Indeed, in females, acute estradiol increases cell proliferation in young but not middle-aged nulliparous rats (12,13). Intriguingly in females, previous reproductive experience (pregnancy and motherhood) can rescue the hippocampus’ response to estrogens later in middle-age, as acute estrogens increased cell proliferation in multiparous rats (13). Thus it is possible that experience, reproductive or otherwise, may restore the ability of androgens to upregulate hippocampal neurogenesis in males in middle age. However, Moser et al. (58) did not see an influence of high fat diet on the androgen modulation of neurogenesis. Intriguingly in our study, the proportion of BrdU/NeuN colabelled cells was affected by age and treatment. DHT increased the proportion of BrdU/NeuN colabelled cells only in middle-aged females, and to our knowledge the first description of such a change in cell fate with aging and DHT. Overall as expected, the proportion of new cells that express mature neuronal protein decreased with age. In middle-aged female mice, letrozole, an aromatase inhibitor blocking the conversion of androgens to estrogens, increases hippocampal neurogenesis (using the immature marker doublecortin; (59)). Together with our findings, this suggests that the hippocampus may still be able to respond to sex steroid hormones in middle age, an effect that varies by sex as in both studies middle-aged females showed increased neurogenesis levels with androgens.

Surprisingly, we found that AR expression increased with aging in both sexes in all hippocampal regions, which to our knowledge has not been reported before. We found ARs were expressed at high and moderate levels in the CA1 and CA3 regions of the hippocampus, respectively, in both young and middle-aged rats. But, consistent with previous research, we did not find expression of ARs in the GCL in young gonadectomized male rats (6,8,60); although there is conflicting evidence which may be due to strain, species, and age differences, as we report AR expression in the GCL at middle age but not in young adults (see below). In Wistar rats, Moghadami et al. (61) and Brännvall et al. (46) found AR expression in the GCL in young gonadectomized and intact male rats. In gonadectomized male mice, ARs are not expressed in the GCL but in males receiving DHT treatment, granule cells did express AR (7). In other rat strains (Bruce-Spruce Long-Evans, Fischer 344 and Sprague Dawley), most studies have found that ARs are not expressed in the GCL (6,8,60) but one study found low to medium AR expression using qualitative analysis in the GCL of intact young male Sprague Dawley rats (62). We also found that DHT increases the expression of ARs in the CA1 in males in line with previous research in young male rats (6) and male mice (7) and in other brain regions (preoptic–hypothalamic regions) (24). In females and males, 2 days of testosterone treatment also showed increased AR expression in the CA1 region (8) but to our knowledge no other studies have investigated AR expression in the female hippocampus after chronic DHT treatment. In female rats, AR expression varies with the estrous cycle (62) as AR expression was highest when estradiol levels are low in the CA1, CA3, and dentate gyrus (62). Furthermore, DHT treatment for 7 days decreased AR expression in most brain regions in intact female rats (62), suggesting that DHT in the presence of gonadal hormones downregulates AR expression in the female hippocampus. In the current study, DHT increased AR expression in gonadectomized females (as well as males), suggesting that gonadal hormones interact to regulate brain levels of AR. As discussed above, previous work has found that androgens increase neurogenesis via the AR. However, in young male rats and mice, ARs are not found in immature neurons (doublecortin expressing cells) in the GCL of the hippocampus (6,7) but instead ARs are expressed in mature neurons (NeuN expressing cells) in the hippocampus (62). We have previously proposed that one mechanism of action is that androgens bind to ARs in the CA3 region and this initiates a retrograde response of a survival factor that targets newborn neurons in the GCL (63). In the current study, we found ARs were expressed in the CA1 and CA3 regions in young and middle-aged male and female rats and their expression increases after DHT treatment in both sexes at both ages. However, we only observed an effect of DHT on new neuron survival in young males. In young females and middle-aged males and females, ARs are expressed in the CA1 and CA3 but binding of DHT to these ARs does not result in the modulation of neurogenesis, possibly indicating different downstream mechanisms of bound AR to modulate neurogenesis in young adult females, and middle-aged males. Interestingly, we found higher expression of AR in the hippocampus of middle-aged compared to young rats of both sexes. In addition, in contrast to young rats, we detected AR-ir cells in the GCL in middle-aged rats. AR mRNA expression also increases in the hippocampus with aging in males (60) but to our knowledge this is the first study showing that hippocampal AR expression increases with aging in females. It may seem paradoxical that although DHT increases neurogenesis via the AR in males (6), despite the increase in AR expression with middle age, DHT no longer increases neurogenesis in middle-aged rats. However, we have previously found that overexpression of the AR in the brain (Nestin-AR) resulted in a failure of DHT to increase neurogenesis in young adult male mice (7). This suggests a ‘dose-response’ of AR expression, with optimal levels needed for DHT to increase neurogenesis. In the preoptic area, the number of AR-ir cells increase with aging in intact male rats (64), but not in castrated males (24), and these are negatively correlated with circulating testosterone levels (64). In young rats, AR expression is dependent on androgen levels and gonadectomy decreases AR expression in the hippocampus (8,60,61). Circulating androgens decrease with age in male rats (65) and therefore we would expect AR expression to decrease with age. Our findings and the ones by Wu et al. (64) are surprising and it is possible that with aging the relationship between androgens and DHT changes. Indeed, we observed that DHT treatment increased AR expression in the CA1 in middle-aged males and females indicating that exogenous regulation of AR expression is similar in both ages. As outlined above, high doses of AR in the brain (in transgenic male mice) result in an abolished neurogenic response to DHT (7). It is possible that the increase in AR expression with aging is responsible for the lack of DHT mediated increase in new neuron survival.

To summarize, our study demonstrates that DHT increases neurogenesis in young adult males but not in young adult females and that aging eliminates the ability of DHT to enhance neurogenesis in males showing sex and age differences in the neurogenic response to DHT. However, in middle-aged rats, DHT treatment increased the proportion of surviving cells expressing the mature neuronal protein (an effect seen in the females only). AR expression in the hippocampus increases with aging and DHT treatment in both sexes, suggesting that ARs do respond to DHT but this does not result in a neurogenic response in females and middle-aged males. Altogether our current and previous research (reviewed in (3)) indicates that androgens and estrogens have sex-specific and age-specific effects on hippocampal neurogenesis.

## Acknowledgments

This work was funded by the Canadian Institutes of Health Research (MOP102568) to LAMG. We thank Dr. Mark Martindale (Whitney Laboratory for Marine Bioscience, University of Florida) for access to an epifluorescent microscope.

